# Longitudinal analysis of visuomotor orientation after optic lobe lesions reveals brain plasticity in *Drosophila*

**DOI:** 10.64898/2026.08.20.745927

**Authors:** Margarida Caio, Dean Rance, Christa Rhiner

**Affiliations:** Champalimaud Foundation, Champalimaud Research, 1400-038 Lisbon, Portugal

**Keywords:** brain injury, stripe fixation, idiosyncratic behavior, *Drosophila*, neural progenitors

## Abstract

Acute brain injury disrupts neuro-glial networks leading to impaired brain function. Although injury induces diverse forms of plasticity, their contributions to brain injury outcome remain poorly understood. We previously showed that targeted stab lesions to the optic lobe (OL) of the adult fly brain induce proliferation of glial and neural progenitor cells. Here, we examined the effect of OL lesions on distinct features of fly behavior, which revealed a specific drop in visual stripe fixation performance acutely after injury, whereas locomotor behavior remained mostly unaffected. Using longitudinal studies of injured individuals, we found that flies significantly regain stripe fixation capacity and idiosyncratic traits one week post injury, suggesting a role for plasticity mechanisms. When the proliferation of adult neural progenitor cells is specifically blocked prior to injury, individuals showed no significant improvements of visual orientation during the identified plasticity window suggesting that progenitor activation may support recovery of stripe approach behavior. Hence the individual tracking of orientation behavior emerges as a suitable quantitative framework for studying functional recovery and inter-individual variability in the adult *Drosophila* brain following brain injury.

## Introduction

Traumatic brain injuries are debilitating, with individuals frequently struggling with diverse cognitive and motor impairments that affect behavior. In human patients, studies typically focus on clinical assessments of sensorimotor function. The current non-invasive imaging techniques provide relatively low resolution across complex brain circuits and are not yet able to monitor plasticity and repair at the cellular level. In turn, studies addressing nervous system injury in model organisms often focus on cellular regeneration, but the impact on behavioral recovery is not examined. Exceptions are well defined spinal cord injury procedures in vertebrates, for which nerve outgrowth, inflammation and glial reactivity are assessed along functional analysis of limb and bladder function (Paramos-de-Carvalho et al., 2021; Nogueira-Rodrigues et al, 2022; Liebscher et al., 2005).

In *Drosophila*, several nerve injury paradigms have elucidated key responses to nervous system damage such as the coordinated breakdown of neurons in Wallerian degeneration via dSarm and Nmnat-mediated signaling (Osterloh et.al., 2012; Neukomm et al., 2014; Avery et al., 2009; Fang & Bonini, 2012) and neuronal debris clearing via Draper-mediated engulfment (MacDonald et al., 2006; Ziegenfuss et al., 2008; Doherty et al., 2009; Logan et al., 2012). In addition, injury to the larval ventral nerve cord (VNC) has revealed dedicated gene networks underlying glial regenerative responses (Harrison et al., 2021; Kato et al., 2017). Also, laser-induced axon ablation in the larval VNC have shown that the glial metabolic state regulates axon regeneration and functional recovery (Li et al., 2020).

Although a significant fraction of research focused on the developmental stage, studies in the adult fly brain also show injury-induced plasticity that support repair. Stab lesions to the central brain or optic lobes (OLs) cause glial reactivity, glial proliferation and the activation of quiescent neural progenitor cells (Kato et al., 2009; Crocker et al., 2021; Simões et al., 2022). More recently, a nerve crush paradigm targeting the adult VNC was described to induce reactive plasticity (Losada-Pérez et al., 2021), including glia-to-glia fate changes (Casas-Tintó et al., 2025). In the setting used, the targeting of different VNC hemisegments could be linked to specific locomotor impairment of fly legs, for which locomotion was scored as either improved or unchanged based on visual inspection of leg dragging. In a model of closed-head injury (head compression), concussions caused temporary ataxia as assessed by the reflex of flies to regain posture when placed on their back and impairments in climbing abilities (Saikumar et al., 2020), which was also observed in a repeated head impact model (Behnke et al., 2021). Moreover, concussion elicited seizure-like twitches in injured flies (Saikumar et al., 2020). Interestingly, flies receiving a central brain puncture showed acutely disrupted locomotion when surveyed with an activity monitor, but completely recovered behavioral patterns by 2 weeks, suggesting that localized brain damage can induce extensive plasticity that may lead to recovery of function (Crocker et al., 2021). Stab lesions to the adult OLs have been shown to reactivate normally quiescent neural progenitors that express the neuroblast marker and HES1-like transcription factor deadpan (*dpn*) (Fernández-Hernández et al., 2013; Li et al., 2020) and upregulate *myc*, which is required for their proliferation (Fernández-Hernández et al., 2013; Simões et al., 2022). However, it is not yet known whether injury to OLs, which process visual information, causes any defects in visually guided behaviors and if so, whether flies can regain function.

In the visual system, Dorsal Cluster Neurons (DCNs) are responsible for coordinated stripe fixation and approach behavior observed in a Buridan arena (Linnewebber et al., 2020), an open-field stage, on which fly locomotion and orientation towards two visual targets is recorded (Colomb et al., 2012). This visual behavior varies both across different isolates of wild-type strains and among individuals with the same genetic background. This variability has been attributed to stochastic differences in neural circuit assembly during development, with flies showing higher DCN branch asymmetry displaying more pronounced stripe fixation behavior (Linnewebber et al., 2020). Also, other behaviors have been reported to exhibit unique traits peculiar to an individual, termed idiosyncrasies, as described for phototaxis (Kain et al., 2012; Honegger & De Bivort, 2018; Krams et al., 2021), locomotor handedness (Ayroles et al., 2015; Buchanan et al., 2015; Skutt-Kakaria et al., 2019), olfactory signature (Honegger et al., 2019; Rihani & Sachse, 2022; Churgin et al., 2023) and social distancing (Bentzur et al., 2020). Idiosyncratic behaviors are also present in other species, but few studies have analyzed recovery of function after nervous system damage and the possible reappearance of idiosyncrasies (Biernaski & Corbett, 2001; Farr and Wishaw, 2002; Mogensen & Malá, 2009).

Here, we interrogated the effect of OL injury on recovery of visuomotor behavior by longitudinal tracking of individuals in injured and control cohorts and uncover a key plasticity window following injury during which regain of stripe approach behavior is observed.

## Results

### Optic lobe injury acutely impairs visually guided orientation

To understand the effect of OL lesions (Fernández-Hernández et al., 2013) on fly behavior, we recorded flies exploring a Buridan arena (Colomb et al., 2012), which allowed to track different parameters of fly locomotion and orientation. In this paradigm, flies explore a circular platform surrounded by water and alternate between fixation of two inaccessible stripes at opposing ends of the arena, creating an innate fixation and anti-fixation response (Fig. 1A). We recorded wild-type flies (*Canton S*) with bilateral, unilateral or no OL lesion, one day and one week post injury (Fig. 1B). Flies with OL lesions did not display differences in diverse parameters assessed for locomotion (activity time, nr of walks, nr of pauses etc.), except for reduced median speed (Fig. S1A and data not shown), but showed significantly impaired stripe fixation capacity shortly after injury, observable by increased strive deviation angles, which was especially pronounced for bilateral OL injury (BI) (BI 1dpi: 35° ± SEM vs 24° ± SEM controls) (Fig. 1C). Remarkably, stripe fixation ability was recovered to a large extent by 1 week post injury (1wpi), suggesting recovery of OL function after the lesion (Fig. 1C).

**Figure 1.**
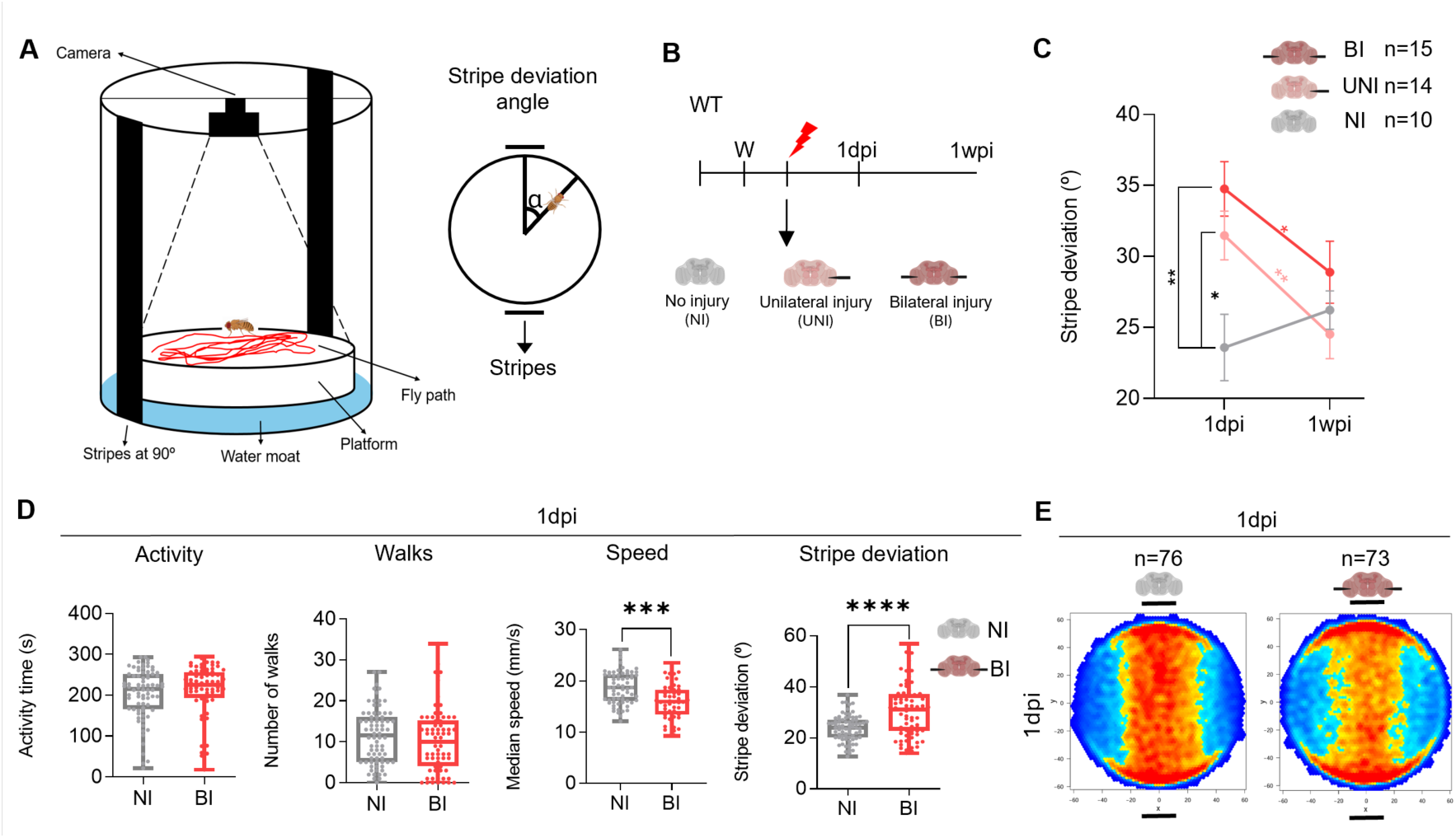
Impaired stripe fixation behavior after optic lobe injury. **A.** Scheme depicting the features of the Buridan arena. Two black stripes are placed at either end as visual hallmarks. Stripe deviation angle indicates deviation from central stripes. **B.** Timeline showing the experimental design. Black filaments indicate unilateral (UNI) or bilateral (BI) stab lesions to the OL lobe of the adult fly brain versus no injury (NI). Wings (w) are clipped 1 day post-hatching to track walking behavior. Orientation behavior is recorded one day post brain injury (1dpi) and 1 week post injury (1wpi). **C.** Graph depicting stripe deviation of flies’ post injury. NI: non-injured (grey), n=10; UNI: unilateral OL injury (light red), n=14; BI: bilateral OL injury (red) n=15. Non-parametric repeated measures two-way ANOVA with Tukey’s test for multiple comparisons. Error bars depict SEM. *p<0.05; ** p<0.01. Black stars represent comparisons between conditions and colored stars between time-points of the same condition. **D.** Quantified locomotor and orientation parameters in the Buridan arena 1 day post injury (1dpi). NI (grey) n=76, BI (red) n=73. Mann–Whitney U test. Error bars depict SD for all. *** p<0.001 and **** p<0.0001. **E.** Transition plots for NI and BI at 1dpi. Transition heatmaps represent the distribution of flies on the platform. Deep red indicates high occupancy (95%), deep blue, very low occupancy.

To further test the causal relation between OL lesion and the ability to approach stripes in a linear manner, we conducted a similar experiment, with a bigger sample size (N ≥ 73) and focused on flies that were bilaterally injured, avoiding compensation by the non-injured eye. We again observed that injured flies showed a significant increase in their stripe deviation angle 1 day post injury (1dpi), consistent with previous results (Figs. 1D and 1E), suggesting that stripe deviation is a parameter critically affected by OL lesions. We further detected a decrease in the median speed of bilaterally injured flies 1dpi, whilst other locomotor parameters stayed unaffected acutely after injury, indicating that the behavioral effect is mostly related to visual impairment (Fig. 1D).

Previous work has established that flies display interindividual variability in their stripe fixation behavior (Brembs & Colomb, 2015; Linnewebber et al., 2020). A population of wild-type flies typically presents a normal distribution, with a part of flies exhibiting highly accurate approach behavior while others show a weaker fixation response. This behavioral individuality arises from stochastic dorsal cluster neuron (DCN) arborization during development, with individuals showing high asymmetry between the two OLs tending to be high responders for stripe fixation (Linnewebber et al., 2020). Indeed, our measurements of large numbers of wild-type flies (Canton S isolate) also showed significant variability in stripe fixation capability across individuals (Fig. 2A). These differences in stripe fixation capability of individuals poses difficulties for the interpretation of data regarding improvement of this behavior post injury.

**Figure 2.**
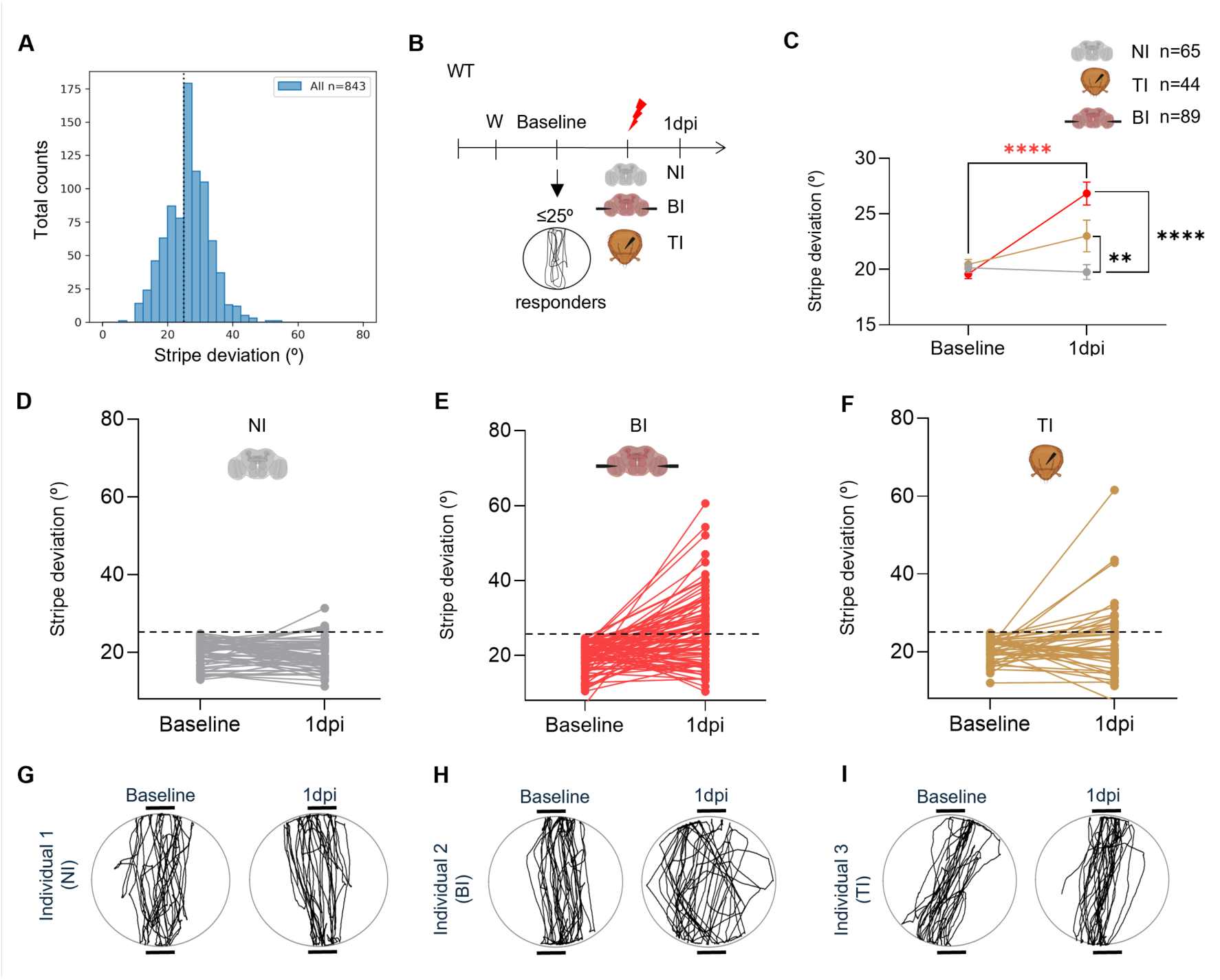
High responders show specific impairments in visual approach behavior. **A.** Histogram showing the distribution of stripe deviation for Canton S flies. The line at 25° marks the chosen threshold for responders. **B.** Scheme depicting the experimental design. Baseline performance was measured for all groups 1 day before injury. NI: non-injured, TI: thorax injury, BI (brain injury, bilateral OL lesion). Only responder flies (stripe deviation ≤ 25°) were injured and assessed one day post injury (1dpi). W: wing clipping. **C.** Graph depicting stripe fixation behavior. NI: non-injured (grey, n=65), BI: bilateral optic lobe lesion (red, n=89), TI: thorax injury (brown, n=44). Non-parametric repeated measures two-way ANOVA with Tukey’s test for multiple comparisons. Error bars depict SEM ** p<0.01; **** p<0.0001. Black stars represent comparisons between conditions and colored stars between time-points of the same condition. **D-F**. Stripe deviation angles for all recorded individuals at baseline and 1dpi, for NI (D), BI (E), TI (F). NI: non-injured (gray, n=65), BI: bilateral optic lobe lesion (red, n=89), TI: thorax injury (brown, n=44). **G-H**. Representative individual walking traces for non-injured flies (NI), flies with brain injury (BI) or thorax injury (TI) at baseline and 1dpi.

In a next set of experiments, we therefore recorded baseline stripe fixation behavior for all flies prior to injury and performed stab lesions only on selected “responder” flies that showed a maximum of 25° deviation angles or lower at baseline (Fig. 2B). In addition, we also recorded orientation behavior of flies with a thorax lesion. In line with previous experiments, the injured stripe-responder individuals showed strongly impaired fixation behavior acutely after injury compared to non-injured controls (Fig. 2C-F).

To understand if the effect was specific to OL lesions, we also performed a thorax puncture. Flies with thoracic injury mostly showed a very mild increase in stripe deviation angle, except for 2 individuals that display significant increases (Fig. 2C) compared to the consistent increases detected in the OL injury group (Figs. 2C and 2E) indicating that loss of stripe fixation behavior is strongly dependent on OL injury. The loss of visual orientation after OL lesions was also directly observable in the recorded walking paths of individuals with OL lesions (Figs. 2G-Other parameters such as the median speed, activity time and number of walks performed by flies were not significantly affected by OL or thorax injury (Fig. S2A). Comparing the results from two large datasets (Fig. 1D vs Fig S2A), the effect of OL injury on median speed was thus inconsistent. Overall, the results identify the stripe deviation angle (visual orientation) as the key parameter that is selectively impaired following OL injury.

### Visual orientation improves during an early plasticity window

As stripe fixation behavior was strongly impaired acutely after brain injury, we next asked if flies showed improvements at delayed time-points, coinciding with previously characterized regenerative responses including the activation of normally quiescent neural progenitor cells (Simões et al., 2022). To address this, we focused on cohorts of flies, in which the OL targeted stab lesion had caused a clear loss of stripe fixation behavior 1 dpi (injury-impaired flies with > 30° stripe deviation angle) compared to their individual baseline performance (Fig. 3A). We next set out to perform longitudinal studies of individual flies with such defined lesions compared to controls by following their performance in the Buridan arena over 2 weeks (Fig. 3B).

**Figure 3.**
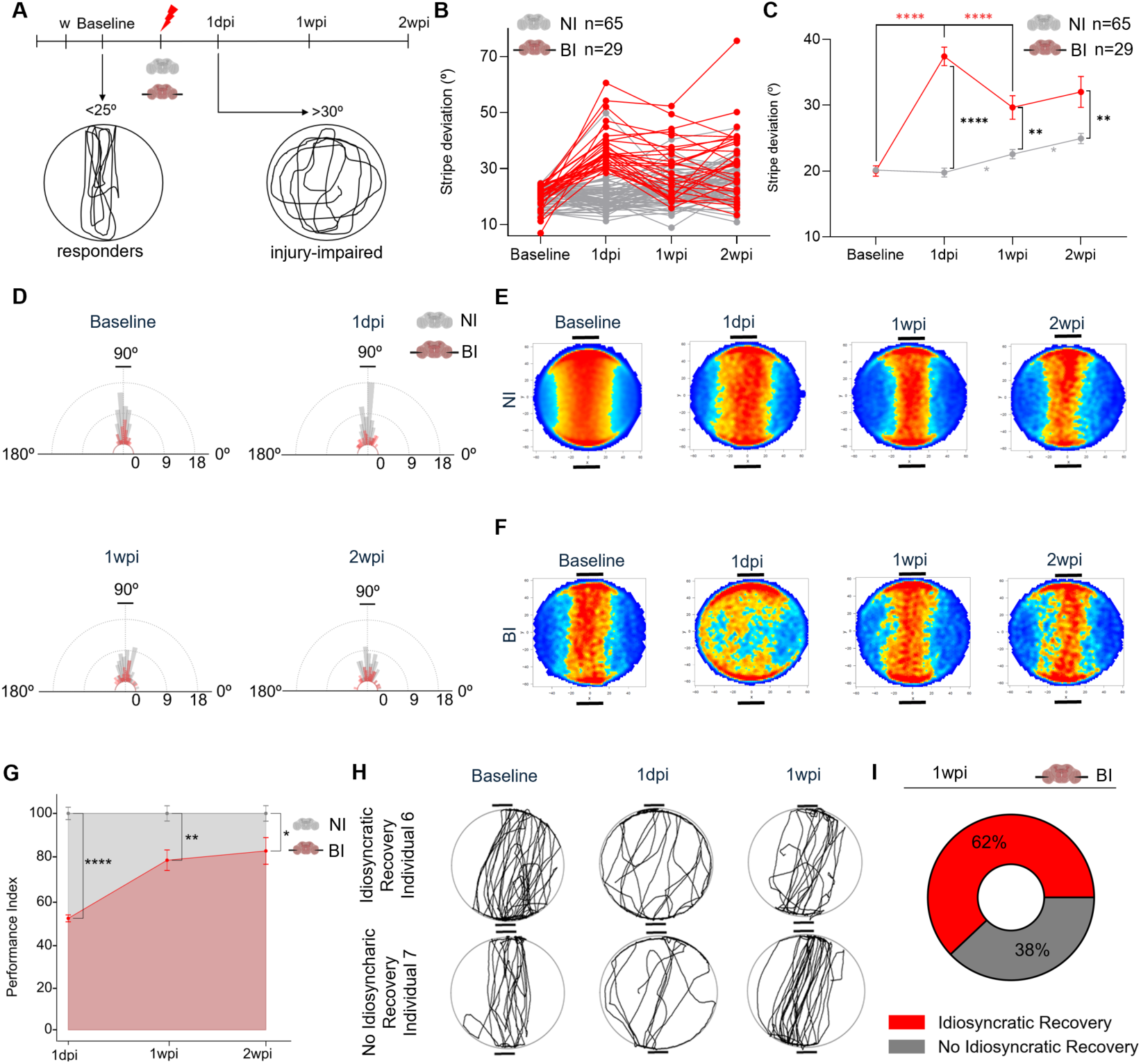
Individual patterns of visuomotor recovery after OL lesions. **A.** Scheme of experimental design. Baseline performance was recorded for all groups 1 day before injury. NI: non-injured, BI (brain injury, bilateral OL lesion). Only responder flies (stripe deviation angle ≤ 25°) were injured and assessed one day post injury (1dpi). Injury-impaired flies (stripe deviation angle >30°) were monitored at 1dpi,1wpi and 2wpi. **B.** Graph depicting stripe fixation behavior for non-injured (NI) individuals (grey, n=65) and bilaterally OL-lesioned wild-type flies (BI) (red, n=29). **C.** Graph showing mean stripe fixation behavior for NI and BI flies. **D.** Polar plots showing the distribution of stripe fixation angles of NI and BI flies. **E, F.** Transition plots showing arena occupancy of control (NI) (E) and OL-injured (BI) populations (F) at baseline, 1dpi and 1wpi. **G.** Performance Index expressing the extent of behavioral recovery in % respective to the average age-matched non-injured control population set to 100% of expected behavior at the respective timepoints at 1dpi, 1wpi and 2wpi. **H.** Representative individual walking traces of flies showing idiosyncratic recovery or no recovery at 1dpi and 1wpi compared to their individual baseline before injury. **I.** Pie chart depicting the percentage of flies that do (red) or fail to recover (grey) idiosyncratic approach behavior (stripe fixation) one week post injury (1wpi).

Interestingly, we observed that flies measured one week after injury showed a significant improvement in visual orientation, reflected in a lower stripe deviation angle (Fig. 3C), with a large fraction of individuals (34%) showing marked improvements compared to their acute impairment 1 day post injury. At a delayed timepoint (2 weeks after injury), the orientation behavior slightly deteriorated rather than showing further recovery. This trend was also observed in the non-injured population, representing a general drop in behavioral performance associated with fly aging (Fig 3C).

The improved navigation one week after injury was also clearly observable at the population level, with a significant fraction regaining precise approach behavior to the displayed black landmarks resulting in predominant walking in a central stripe of the arena compared to non-injured controls, which maintained central arena occupation over the 2-weeks period (Figs. 3D-3F). When compared against realistic recovery that can be expected of age-matched non-injured controls, wild-type flies regained almost 80% of stripe fixation capacity (Fig. 3G), which did not notably improve in the following week, reinforcing the existence of a crucial 1-week plasticity window following injury.

### Individuality of orientation behavior

It has been previously reported that flies show individual biases in stripe fixation in the arena, with some individuals exhibiting more left-leaning or right-leaning approach behavior (Linnewebber et al., 2020). This feature allows studying if such individual traits are regained after injury in addition to recovering the “standard” responder behavior. We analyzed the individual walking traces (Fig. 3H) by implementing a method to statistically track the preferred angle (code available in methods) and found that by one week after the injury, 62% of the injured individuals showed recovery of their idiosyncratic stripe approach behavior, whereas 38% did not recover such individual traits displayed prior to injury (left or right) (Figs. 3H and 3I). The observed recovery is remarkable and highlights the possible involvement of injury-induced plasticity, which allows to restore particular visuomotor patterns in a significant number of individuals.

### Inhibition of progenitor activation restricts recovery of stripe deviation

Previous work has demonstrated that an OL stab lesion triggers adult neural progenitors to exit quiescence and proliferate (Fernández-Hernández et al., 2013; Simões et al., 2022). These rare progenitors appear to be scattered throughout the medulla cortex and express the HES1-like transcription factor *dpn* (Fernández-Hernández et al., 2013; Li et al., 2020). To understand whether the observed recovery in stripe fixation behavior is dependent on progenitor activation, which upon cell cycle entry can give rise to new neurons at the lesion site, we specifically suppressed *myc* in adult neural progenitor cells with *dpnT2A; tubGal80^ts^ using myc RNAi,* which efficiently blocks progenitor proliferation (Simões et al., 2022). RNAi was activated in adult flies with a temperature shift 3 days prior to injury, leading to inactivation of the thermosensitive (ts) Gal4 suppressor, Gal80^ts^ (Fig. 4A). When recording baseline stripe fixation behavior in this genetic background, we encountered a 50-63% reduction in responder frequency compared to Canton S, which implied pre-screening a high number of animals.

**Figure 4.**
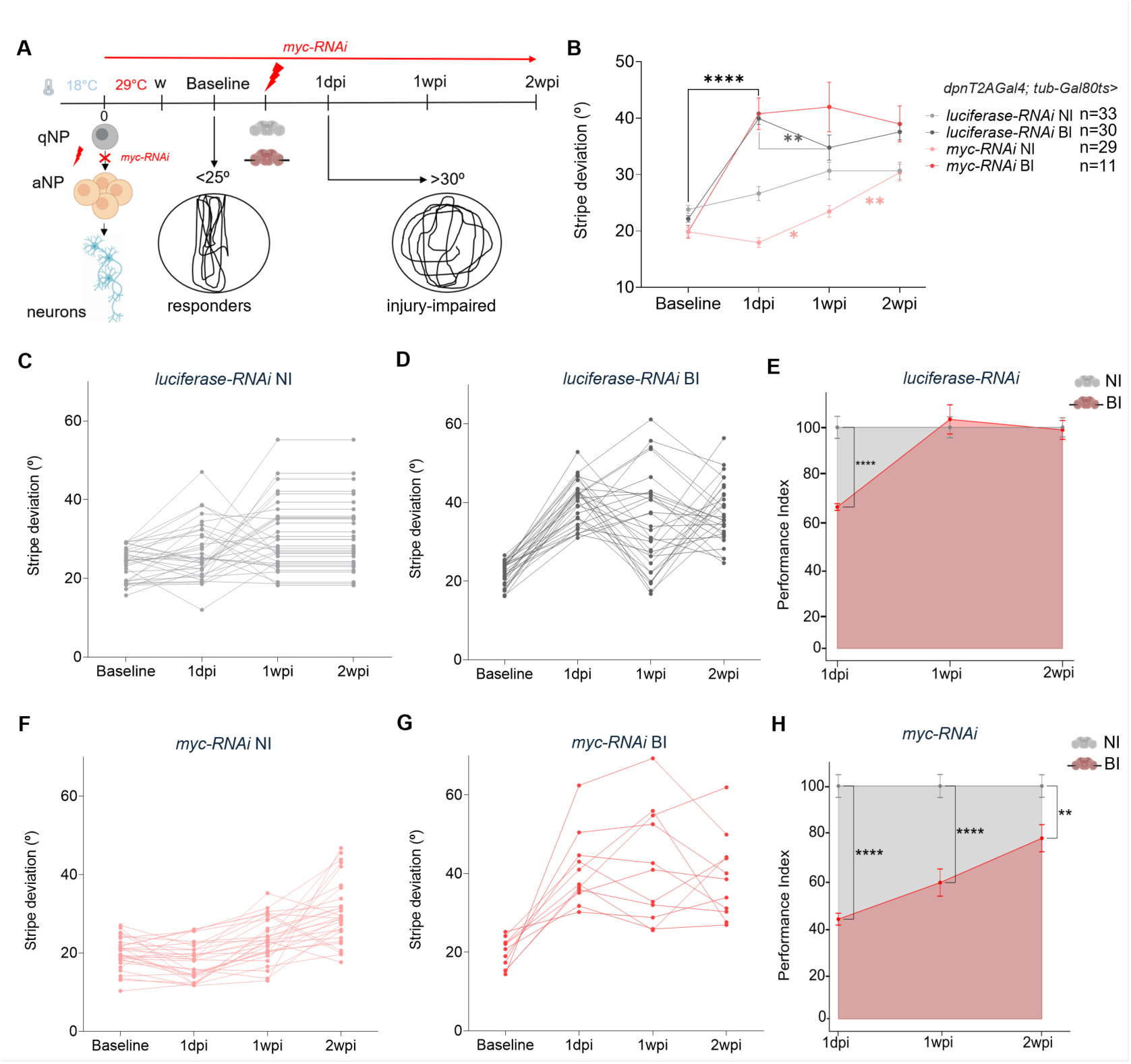
Suppression of neural progenitor activation results in reduced recovery parameters. **A.** Experimental design. Flies were reared at 18°C until hatching, RNAi is activated 3 days prior to injury by shifting flies to 29°C for optimal RNAi activation. Expression of *myc*-RNAi in neural progenitors is driven by *dpnT2Agal4; tub-Gal80ts*, preventing their activation in response to injury. *Luciferase-RNAi* is used as control for RNAi activation. **B.** Graph showing stripe deviation angles (in degrees). *Luciferase-RNAi* NI (light grey) n=33, *luciferase-RNAi* BI (dark grey) n=29, *myc-RNAi* NI (red) n=29, *myc*-*RNAi* BI (light red) n=11. Non-parametric repeated measures two-way ANOVA with Tukey’s test for multiple comparisons. Error bars depict SEM, * p<0.05; ** p<0.01; **** p<0.0001. Black stars represent comparisons between conditions and colored stars between time-points of the same condition. **C, D.** Graph depicting stripe fixation behavior for individual NI (n=33) (C) and injured (BI) *luciferase-RNAi* flies (D) (n=30). **E** Performance Index expressing the extent of behavioral recovery in % respective to the average age-matched non-injured control population (*dpnT2A, tub-Gal80ts>luciferase RNAi*) set to 100% of expected behavior at the respective timepoints at 1dpi, 1wpi and 2wpi. **F,G.** Graph depicting stripe fixation behavior for individual NI (n=29) (F) and injured (BI) *myc-RNAi* flies (F) (n=11). **H.** Performance Index expressing the extent of behavioral recovery in % respective to the average age-matched non-injured control population (*dpnT2A, tub-Gal80ts>luciferase RNAi*) set to 100% of expected behavior at the respective timepoints at 1dpi, 1wpi and 2wpi.

Nevertheless, we were able to record data for smaller cohorts with activation of RNAi in adult neuronal progenitors. Consistent with previous findings, the injury caused a significant impairment of orientation acutely after injury both in *myc* RNAi and control flies (*luciferase* RNAi), compared to the respective non-injured lines. The trajectory of flies with intact *myc* activation showed a trend to improved stripe fixation behavior although the regain in performance was not pronounced as in wild-type (Canton S) flies (Fig. 4B). In contrast, stripe deviation angles remained high in flies with Myc suppression in adult progenitors (no improvement of orientation) when retested 1 week after injury (Fig. 4B). Although limited to a smaller number of analyzed lesions, the plotting of the individual recovery curves (Figs. 4C-4H) revealed a significant fraction of control flies (*luciferase* RNAi) showing clear trajectories of recovery (Fig. 4D) and reaching performance of age-matched non-injured controls 1wpi (Figs. 4C and 4E), the flies with suppressed progenitor activation displayed mostly flat recovery trajectories (Fig. 4G) reaching less than 60% performance of behavior of age-matched, non-injured flies with myc RNAi activation (Fig. 4H).

Altogether, these results suggest that progenitor activation contributes to recovery during an early plasticity window after brain injury, whereas the inhibition prevents or significantly delays restoration.

## Discussion

### Functional Impact of Optic Lobe Lesions and Behavioral Recovery

Our results demonstrate that targeted stab lesions to the adult *Drosophila* optic lobe (OL) specifically impair visually guided orientation, leaving locomotor ability largely intact. While previous models utilizing high-impact trauma or head compression reported long-lasting deficits in climbing, walking, and spontaneous seizures (Saikumar et al., 2020; Behnke et al., 2021), our OL stab lesion paradigm, by causing more localized damage, allows to observe a partial recovery. which is supported by pronounced injury-induced plasticity in the OL. Flies subjected to bilateral OL lesions exhibited a significant increase in stripe deviation angles acutely at 1 dpi, identifying this parameter as a sensitive metric for visually-guided orientation. Remarkably, injured wild-type flies regained up to 80% of their orientation performance within one week (Fig. 3G). This temporal window of recovery coincides with the previously observed emergence of newly formed differentiated neurons in the OL following division of adult neural progenitor cells (Fernández-Hernández et al., 2013). The recovery observed supports the notion that the adult fly brain shows plasticity in response to injury which improves brain injury outcome at the behavioral level.

In particular, we the results suggest that the regaining of vision-driven orientation is aided by activation of neural progenitors. Unlike control flies, individuals with inhibited *myc* expression in progenitors showed no improvement in stripe fixation during the plasticity window, although the results were less pronounced than with wild-type flies. The difference is also arising due to the fact that the monitored flies under RNAi conditions were older due to the 3-day RNAi activation period and the higher temperature used (29°C for optimal RNAi induction vs 25 °C for wild-type flies in Fig. 3A), highlighting that any deviation from the early 1-week plasticity window can restrict recovery.

Interestingly, the findings suggest that the improvement of orientation is at least partly dependent on adult neural progenitor activation and possible integration of new neurons apart from other envisageable plasticity (recruitment of other circuits, synaptic adaptation). Our results therefore highlight that injury-induced plasticity and progenitor proliferation play an important role in restoring complex visual function in the adult brain.

### Inter-individual Variability and Idiosyncratic Recovery

In agreement with the findings of Linneweber et al. (2020), our population exhibited significant inter-individual variability in baseline stripe fixation and trajectory biases. This behavioral "personality" is due to the stochastic wiring and branching asymmetry of the DCN neurons established during development. We interrogated whether individuals would restore not just the general orientation behavior, but the individual’s unique pre-injury bias, possibly due to injury-induced repair, although we cannot exclude that inflammation may also play a role in the acute impairment. While 62 % of injured individuals recovered these their stripe approach traits, others regained only the ability to orient toward the stripes without reinstating their idiosyncratic pre-injury behavior. This partial restoration is still remarkable, given the restricted environment of the adult brain regarding plasticity. Overall, complete recovery after nervous system damage is rarely observed, with individuals regaining on average 70% of their pre-injury abilities (Prabhakaran et al., 2007; Winters et al., 2014). Still, variability and the use of alternative plasticity mechanism and behavioral adaptation have been suggested to modulate an individual behavior and ultimately ensures species survival under stress (Kain & Bivort, 2015; Werkhoven, 2019; Krams, 2021).

Stab lesions have also been shown to induce rapid glial divisions in the adult fly brain and changes in membrane extensions (Kato et al. 2017, Simões et al., 2022). Therefore, glial state changes and increased glial support may also modulate brain restorative processes.

Altogether, this study establishes that flies suffering localized damage to the optic lobes show significant recovery in stripe fixation behavior, which is selectively impaired following OL lesions. By quantitatively tracking individuals in the Buridan arena, we demonstrate individual recovery trajectories during a key plasticity window and which are partly modulated by injury-induced plasticity, in particular progenitor activation. These findings underscore the potential of optic lobe lesions in Drosophila as a promising model to further bridge the gap between altered cellular interactions and recovery of brain functions after acute injury in the future.

## Acknowledgments

We thank the CF fly platform for technical support and the Bloomington and Vienna *Drosophila* stock centers for fly lines. This work has been supported by grant HR23-00860 from LaCaixa & Fundação para a Ciência e Tecnologia (FCT), the ERC-Portugal program (FCT) to C.R. and the Champalimaud Foundation.

## Author contributions

This study was conceptualized by C.R. with contributions from M.C.. Experiments were performed by M.C. and formal data analyses by M.C. and D.R with codes established by D.R. The manuscript was written by M.C. and C.R. with contributions from D.R.

## Disclosure and competing interest statement

The authors declare no competing interests.

## Materials and Methods

### Fly strains

*Drosophila* were reared on food made with these ingredients (3.5L): 280g molasses (barley malt), 70 g Beet syrup, 280 g corn flour, 63 g granulated yeast, 35 g Soy flour, 27g Agar-Agar, 3700ml boiling water. Propionic Acid (28ml) and 15% Nipagin (42ml) are added after cool down to 60°C. Canton S flies were raised at 25°C. Crosses including *tub-Gal80^ts^* (*dpn-T2A-Gal4; tub-Gal80ts/UAS-RNAi-dmyc* (Bloomington, #51454) and *dpn-T2A-Gal4; tub-Gal80ts/UAS-RNAi-luciferase* were raised and kept at 18°C up to 2 days after eclosion and then shifted to 29°C for activation of RNAi. Flies were kept in a 25°C incubator until the behavioral assays and throughout the whole experiment, wing clipping was performed at RT.

### Wing clipping

The day prior to the behavioral recordings in the Buridan arena, flies were anesthetized with CO_2_ and their wings were shortened with precision micro-scissors (FST) to one third of their original length as previously described (Colomb et al., 2012). After wing clipping, flies were left to recover at room temperature with the vial placed on the side until they recovered from anesthesia. Flies were kept in the 25°C incubator until the behavioral assay. In the case of RNAi lines, the wing clipping protocol was done after the 3 days of activation at 29°C, with flies being 5 days old.

### Optic lobe injury

Two-day old adult wild-type flies (Canton S) were anesthetized with CO_2_. A thin metal filament (0.2 mm diameter) was sterilized with 70% ethanol and introduced through both eyes (bilateral injury) or the right eye of adult female flies to the level of the medulla as previously described (Fernandez-Hernandez et.al., 2013). A sterile needle was used to prick the thorax of flies. Non-injured flies were anesthetized for a comparable period. Flies were left to recover at room temperature with the vial placed flat to avoid sticking to food. In the case of RNAi lines, the optic lobe injury protocol was done with flies being 7 days old.

### Stripe Fixation Behavior

Flies were recorded in the Buridan arena (Colomb et al., 2012) at baseline (pre-injury), 1 day, 1 week and 2 weeks post-injury. Age-matched non-injured control flies were assayed in the same days.

The walking activity of each individual fly was recorded for 5 minutes at each time-point, using the Buritrack software (http://buridan.sourceforge.net). Individual tests were re-initialized when flies jumped from the platform or exhibited grooming behavior. Flies were excluded from the trial if they jumped more than 3 times or showed prolonged immobility after placing them on the platform. Flies which appeared to have suffered damage other than the optic lobe lesion during handling were discarded.

Flies that did not show a pronounced stripe fixation behavior during baseline measurements (>25°) were excluded from the experiments analyzing long-term recovery. Fly behavior was analyzed with the CeTrAn software V4 (https://github.com/jcolomb/CeTrAn/releases/tag/v.4) (Colomb et al., 2012). Behavioral parameters analyzed were median speed, distance traveled per minute, turning angle, meander, stripe deviation, activity time, activity bouts and centrophobism index while moving or sitting. Histograms to visualize the range of stripe deviation behavior were analyzed using Python, using the code available in GitHub (https://github.com/margaridasilvacaio-svg/Stripe-Fixation-Analysis).

### Performance Index

The performance index (PI) was calculated using the formula PI (%) *= 100 x (1/*SD*) x 1/(average(1/SD; NI)* where SD is stripe deviation and average (X; inj) is the average of variable X over injury group inj., with this form chosen due to Jensen’s Inequality thereby normalizing the performance to the non-injured age-matched controls.

### Recovery of individual biases

Flies walking trajectories were analyzed for preferred direction of the trajectory – left or right. Flies that recovered their preferred bias were considered as idiosyncratic recovery. Analysis of the trajectory bias were done in Python, using code available in GitHub (https://github.com/margaridasilvacaio-svg).

### Statistical analysis

Statistical tests were applied in accordance with normal or non-parametric data distribution using GraphPad Prism 8 (https://www.graphpad.com/scientific-software/prism/). Non-parametric repeated measures two-way ANOVA with Tukey’s test for multiple comparisons was used for longitudinal follow-ups of individuals. Error bars are shown as ±SD in boxplots and ± SEM in spaghetti plots. ****p < 0.0001, ***p < 0.001, **p < 0.01, *p < 0.05.

---

**Figure S1:**
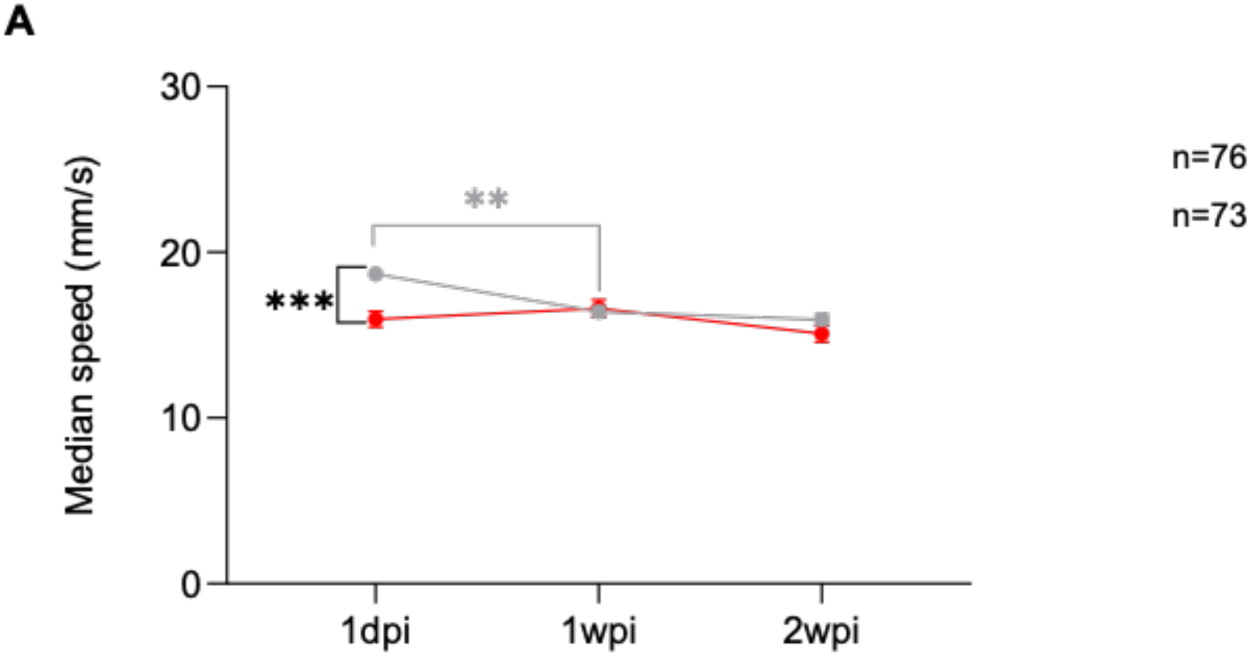
Evolution of median speed parameter over time. **A.** Quantified median speed 1 day post injury (1dpi), one week post injury (1wpi) and two weeks post injury (2wpi). NI (grey) n=76, BI (red) n=73. Non-parametric repeated measures two-way ANOVA with Tukey’s test for multiple comparisons. Error bars depict SEM. **p<0.01, ***p<0.001. Black stars represent comparisons between conditions and colored stars between time-points of the same condition.

**Figure S2:**
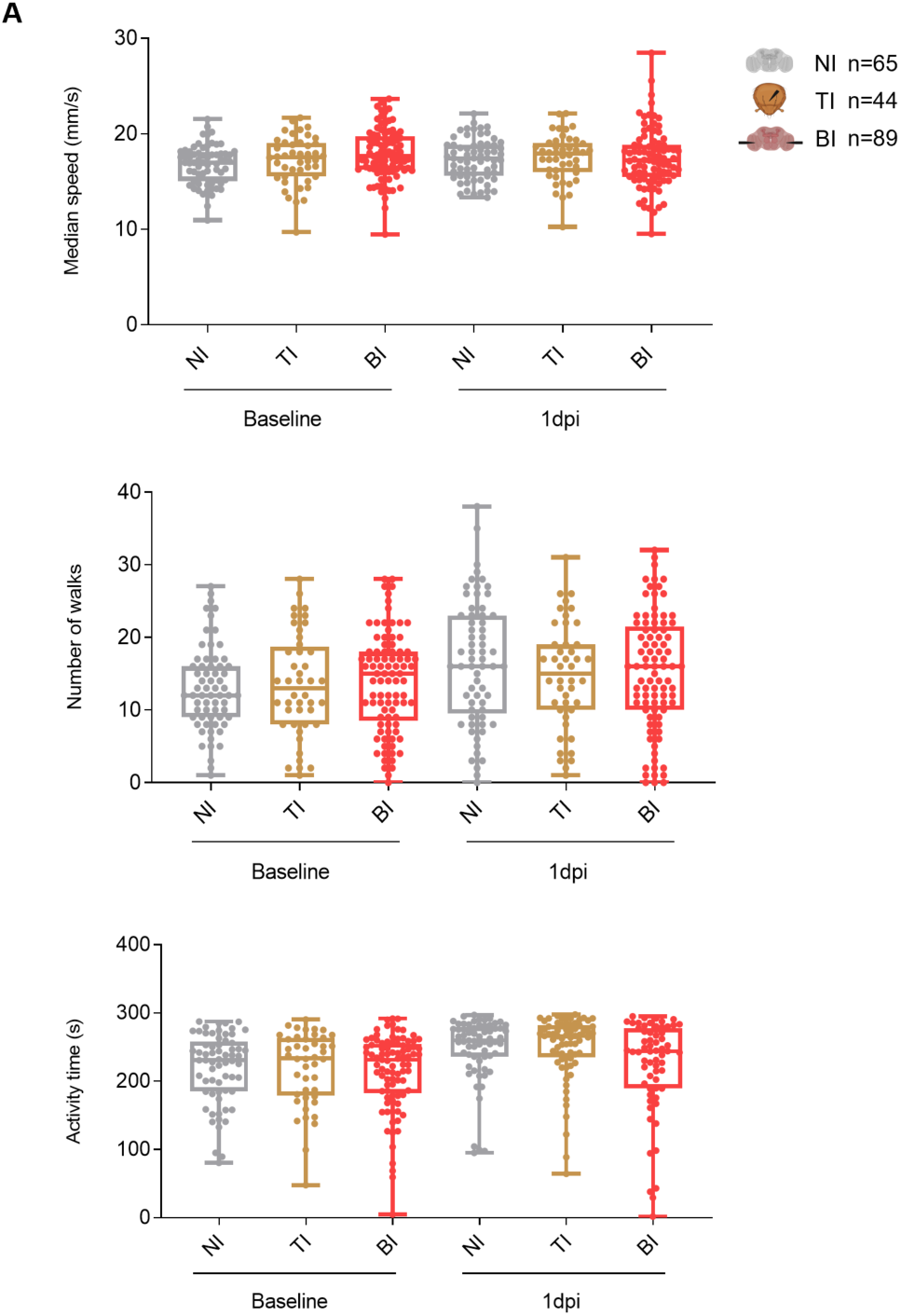
Minor effects of injury on locomotion. **A.** Quantified median speed, nr of walks and activity time at baseline and 1 day post injury (1dpi). NI: non-injured (grey, n=65), BI: bilateral optic lobe lesion (red, n=89), TI: thorax injury (brown, n=44). Applying mixed effect model analyses with Sidak’s correction for multiple comparisons. No significant changes were found.

## Notes

https://github.com/margaridasilvacaio-svg

### Competing Interest Statement

The authors have declared no competing interest.

